# Task errors contribute to implicit remapping in sensorimotor adaptation

**DOI:** 10.1101/263988

**Authors:** Li-Ann Leow, Welber Marinovic, Aymar de Rugy, Timothy J Carroll

## Abstract

Perturbations of sensory feedback evoke sensory prediction errors (discrepancies between predicted and actual sensory outcomes of movements), and reward prediction errors (discrepancies between predicted rewards and actual rewards). Sensory prediction errors result in obligatory remapping of the relationship between motor commands and predicted sensory outcomes. The role of reward prediction errors in sensorimotor adaptation is less clear. When moving towards a target, we expect to obtain the reward of hitting the target, and so we experience a reward prediction error if the perturbation causes us to miss it. These discrepancies between desired task outcomes and actual task outcomes, or “task errors”, are thought to drive the use of strategic processes to restore success, although their role is not fully understood. Here, we investigated the role of task errors in sensorimotor adaptation: during target-reaching, we either removed task errors by moving the target mid-movement to align with cursor feedback of hand position, or enforced task error by moving the target away from the cursor feedback of hand position. Removing task errors not only reduced the rate and extent of adaptation during exposure to the perturbation, but also reduced the amount of post-adaptation implicit remapping. Hence, task errors contribute to implicit remapping resulting from sensory prediction errors. This suggests that the system which implicitly acquires new sensorimotor maps via exposure to sensory prediction errors is also sensitive to reward prediction errors.

## Introduction

Successful goal-directed movement requires the capacity to adapt movements to unexpected changes in the properties of our world or our moving bodies. Perturbations evoke discrepancies between predicted and actual sensory outcomes of movements (i.e., sensory prediction errors), which result in an obligatory, implicit remapping of the input-output relationship between motor commands and the resultant sensory outcomes. Perturbations also elicit either an unexpected failure to attain the reward of hitting the target (i.e., a negative reward prediction error) (Izawa & Shadmehr, 2011), and/or an unexpected punishment of missing the target (i.e., a positive punishment error). It is clear that behavioural responses to perturbations are affected not only by sensory prediction errors, but also by reward prediction errors (Izawa & Shadmehr, 2011; Cashaback *et al.*, 2017; Palidis *et al.*, 2018). However, how reward prediction errors and rewards affect sensorimotor adaptation is not fully understood. Mounting evidence shows that extrinsic rewards and punishments such as gaining or losing pleasing feedback, points, money, or food can modulate sensorimotor adaptation (e.g., Madelain *et al.*, 2011; Galea *et al.*, 2015; Nikooyan & Ahmed, 2015; Gajda *et al.*, 2016; van der Kooij & Overvliet, 2016; Kojima & Soetedjo, 2017; Quattrocchi *et al.*, 2017; Song & Smiley-Oyen, 2017), but less is known about how adaptation is affected by intrinsic rewards associated with accomplishing task goals (Widmer *et al.*, 2016; Kim *et al.*, 2017b).

The unexpected failure to attain the task goal of hitting the target, or task errors, are thought to drive the use of explicit compensatory strategies (Taylor & Ivry, 2011), such as re-aiming to one side of a target when visual feedback of the moving limb is laterally perturbed (Welch, 1969). This explicit learning is thought to be flexible: it can be volitionally disengaged when no longer useful (McDougle *et al.*, 2016). In contrast, sensory prediction errors are known to result in *implicit* remapping: a change in the perceived sensory consequences of motor command (Mazzoni & Krakauer, 2006). Although adaptive behaviour to compensate for perturbations can be driven by sensory prediction errors or reward prediction errors (Izawa & Shadmehr, 2011; Nikooyan & Ahmed, 2015; Cashaback *et al.*, 2017; Palidis *et al.*, 2018), it has been suggested that only sensory prediction errors can produce a change in the system that predicts sensory consequences of motor commands: reward prediction errors alone are insufficient (Izawa & Shadmehr, 2011; Nikooyan & Ahmed, 2015). However, because these studies never made both sensory prediction errors and reward prediction errors concurrently available in the same conditions, it remains unclear whether reward prediction errors modulate implicit remapping resulting from sensory prediction errors.

Here, we tested whether reward prediction errors contribute to implicit remapping resulting from sensory prediction errors. Exploiting the fact that the reward that is intrinsic to typical sensorimotor adaptation tasks is target acquisition, we examined how adaptation was affected by success or failure in acquiring a target (i.e., task errors). When participants were exposed to a 30° rotation of cursor feedback that represented their hand position, we either (1) removed task error by shifting the target mid-movement to align with the (measured) initial cursor direction, such that the cursor always hit the target, (2) enforced task error by randomly shifting the target away from the cursor by 20-30° mid-movement, such that the cursor never hit the target, or (3) allowed standard task error by maintaining a constant target position during the trial. Removing task error dramatically reduced the rate and extent of error compensation to the cursor rotation, but also reduced the amount of implicit remapping resulting from exposure to sensory prediction errors. Enforcing task errors resulted in slower error compensation than the standard task error condition, but did not alter the amount of post-adaptation implicit remapping compared to the standard task error condition. These results suggest that the reward prediction error of hitting or missing targets contributes to the formation of new sensorimotor maps that result from exposure to sensory prediction errors.

## Methods and Materials

### Participants

There were a total of 126 participants (67 female, age range 17-39 years, mean age 21.4+/-0.3. All participants were naïve to visuomotor rotation and force-field adaptation tasks, and were naïve to the aims of the study. Participants received course credit or monetary reimbursement upon study completion. The study was approved by the Human Research Ethics Committee at The University of Queensland. All participants provided written informed consent. This study conforms with the Declaration of Helsinki.

### Apparatus

Participants completed the task using the vBOT planar robotic manipulandum, which has a low-mass, two-link carbon fibre arm and measures position with optical encoders sampled at 1,000 Hz (Howard *et al.*, 2009). Participants were seated on a height-adjustable chair at their ideal height for viewing the screen for the duration of the experiment. Visual feedback was presented on a horizontal plane on a 27” LCD computer monitor (ASUS, VG278H, set at 60Hz refresh rate) mounted above the vBOT and projected to the participant via a mirror in a darkened room, preventing direct vision of her/his hand. The mirror allowed the visual feedback of the target (a 0.5 cm radius yellow circle), the start (a 0.5 cm radius white circle), and hand cursor (0.5 cm red radius) to be presented in the plane of movement, with a black background. The start was aligned approximately 10cm to the right of the participant’s mid-sagittal plane at approximately mid-sternum level. An air-sled was used to support the weight of participants’ right forearms, to minimize possible effects of fatigue.

### General Trial Structure

Participants made centre-out reaching movements while grasping the robot arm. Targets appeared in random order at one of eight locations at a radius of 9 cm from a central start circle. The target locations were distributed uniformly throughout 360° (0°, 45°…. 315°). At the start of each trial, the central start circle was displayed. If participants failed to move the hand to within 1cm of the start circle after 1 second, the robotic manipulandum passively moved the participant’s hand to the start circle (using a simulated 2 dimensional spring with the spring constant magnitude increasing linearly over time). A trial was initiated when the cursor remained within the home location at a speed below 0.1cm/s for 200ms. We used a classical timed-response paradigm (e.g., Schouten & Bekker, 1967) to manipulate movement preparation time (Leow *et al.*, 2017). Across all conditions, a sequence of three tones, spaced 500ms apart, was presented at a clearly audible volume via external speakers. Participants were instructed to time the onset of their movements with the onset of the third tone, which was more highly-pitched than the two previous, and slice through the target with their cursor. Movement initiation time was identified online as when hand speed exceeded 2cm/s. Targets appeared at 1000ms minus a display latency (27.6 ± 1.8ms), before the third tone. Thus, target direction information became available 972ms before the desired initiation time. When movements were initiated 50ms later than the third tone, the trial was aborted: the screen went black and the text “Too Late” was displayed on the feedback screen. When movements were initiated more than 100ms before the desired initiation time, the trial was aborted: the screen went black and a “Too Soon” error message was displayed. Thus, movements had to be initiated between 872 and 1022ms of target presentation. No visual feedback about movements was available when trials were aborted, and so such trials were immediately repeated.

Across all conditions, sensory prediction errors were imposed via a 30° rotation of cursor feedback representing the hand position. Half of all participants encountered a clockwise 30° rotation and half encountered a 30° counterclockwise rotation. Task errors were manipulated in three conditions. In the **StandardTaskError** condition, the target remained stationary throughout the trial, such that whether or not the perturbation evoked a task error was contingent on the participant’s reach direction. Task errors were removed in the **NoTaskError** condition by moving the target to align with the direction of cursor velocity when the hand had moved 4cm (of the 9cm distance) from the start position. This is analogous to moving a basketball hoop towards the ball mid-flight; the ball always goes through the hoop regardless of the person’s actions. Finally, in the **EnforcedTaskError** condition, task errors were enforced on every trial, but were uninformative: the target was shifted randomly by 20° to 30° clockwise or counterclockwise from the cursor direction when the hand had moved 4cm from the start. This is analogous to moving a basketball hoop away from the ball’s trajectory; participants can never get the ball through the hoop regardless of where they shoot. Across all conditions, cursor feedback was displayed after the hand had moved 4cm from the origin (i.e., the point at which cursor direction was measured to define potential target shifts). All participants completed the following blocks, where 1 cycle contained 1 trial to each of the 8 targets (target order was random within each cycle). **Baseline** (6 cycles): no rotation of visual feedback. **Adaptation** (60 cycles): 30° rotation of visual feedback. **No feedback** (6 cycles): Upon leaving the start circle, no feedback about movements were available. Before this block, participants received explicit instructions that the computer no longer imposed any disturbance to the visual feedback, and that they should aim straight towards the target. Between each block, there was a small delay to allow loading of the computer code for different experimental blocks and/or experimental instructions. The task error manipulations were employed only during the adaptation block.

For the main experiment, we ran three participant groups (one for each of the three experimental conditions: StandardTaskErrors, n=30, 23 female, mean age 20.5, range: 17-34 years, NoTaskErrors, n=32, 19 female, mean age 21, range 17-39, and EnforcedTaskErrors, n=32, 23 female, mean age 21.4, range 18-33). We also ran a follow-up study with 30 instead of 60 adaptation cycles, with two participant groups (StandardTaskError, n=16, age 18-22, 11 female; NoTaskError, n=16, age 18-30, 10 female).

### Data analysis

Movement onset time was taken as the time at which hand speed first exceeded 2 cm/s. Movement direction was quantified at 20 percent of the movement distance. This procedure ensured that movement direction was quantified at less than 200ms into the movement, at which time the size of online corrections in hand position is small. Reaches with absolute initial direction errors greater than 60° with respect to the target (movements that were more than 60° to the left or the right of the target) were considered outliers, and were removed from analyses (EnforcedTaskError, 0.66%, NoTaskError, 0.84%, StandardTaskError, 1.45%). Excluding these trials did not have any qualitative impact on the results. Trials were averaged in cycles of eight (one trial for each target angle) for statistical analysis. Reach direction errors for participants who experienced counterclockwise rotations (-30°) were sign-transformed for combined analysis with data for participants who experienced clockwise (+30°) rotations.

Intrinsic biases in reaching direction can affect adaptation (Ghilardi *et al.*, 1995; Vindras & Viviani, 1998; Morehead & Ivry, 2015). Intrinsic biases were evident in the baseline block, as reaches deviated significantly from 0 in the last baseline cycle (p =.001). To estimate intrinsic biases, we averaged reach directions from baseline cycles 2 to 6, and then subtracted this value from all cycles in all adaptation, no-feedback, and washout cycles, similar to previous work (Leow *et al.*, 2017). All subsequent analyses were run on bias-corrected reach directions.

We tested how the different task error conditions altered the time-course of **adaptation** by running mixed ANOVAs with the within-subjects factor Cycle (reflecting changes in reach direction across increasing cycles) and the between-subjects factor Condition (StandardTaskErrors, ConstantTaskErrors, and NoTaskErrors) for the first 30 adaptation cycles. Partial eta-squares were used to report ANOVA effect sizes, with values in excess of 0.14 considered large. When Mauchly’s test of sphericity was statistically significant, the Greenhouse-Geisser correction was used to adjust degrees of freedom.

To test the completeness of adaptation, we estimated the **asymptote** by taking the mean of the last 5 cycles (adaptation cycle 56 to 60). **Disengagement of explicit learning** after notification of the perturbation removal was estimated as the difference between the first no-feedback cycle and the last adaptation cycle. **Implicit remapping** was estimated as the mean of the first no-feedback cycle after notification of the perturbation removal. To test if these measures differed between experimental conditions, we used one-way ANOVAs and follow-up t-tests and Cohen’s d to estimate effect sizes when Shapiro-Wilk tests showed no violations of normality. Cohen’s d values of .8, .5, and .2 represented large, medium, and small effect sizes. When Shapiro-Wilk tests showed violations of normality, we used Kruskal-Wallis tests, followed by Mann-Whitney U-tests, with effect sizes quantified as *r* (Fritz *et al.*, 2012). For *r*, a large effect is .5, a medium effect is .3, and a small effect is .1 (Fritz *et al.*, 2012). Bonferroni corrections were applied in the cases of multiple comparisons. Only two-sided tests were used.

Statistical analyses were performed with JASP (Version 0.8.5) and SPSS. An alpha level of .05 was used. Graphs were plotted with GraphPad Prism version 7.00 for Windows, GraphPad Software, La Jolla California USA, www.graphpad.com.

## Results

### Removing task error reduced error compensation

We examined how manipulating task errors altered how people adapted reaching movements to sensory prediction errors evoked by rotating cursor feedback of hand position. Before encountering the cursor rotation, participants showed a counterclockwise reach bias, although this bias did not differ reliably between groups. Figure 1a shows bias-corrected and cycle-averaged movement directions for each task error condition. Values closer to the ideally adapted movement direction (-30°) indicate more complete compensation for the cursor rotation.

**Figure 1a.**
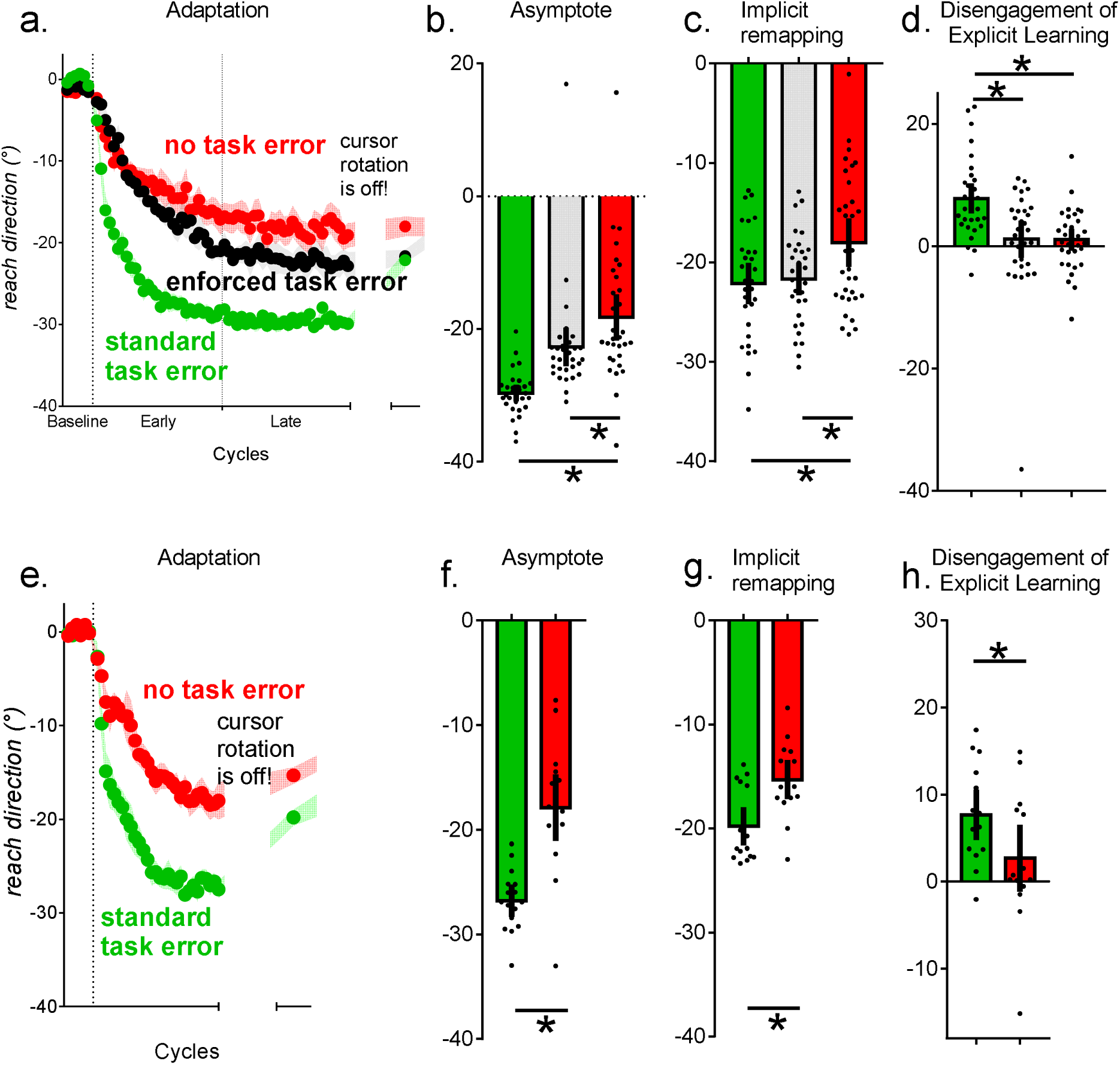
Group mean+/-SEM for reach directions across all cycles (1 cycle=8 trials=1 visit to each of the 8 targets). Reaches closer to −30° represent more complete error reduction. Adaptation was slowest and least complete with NoTaskErrors (red), followed by EnforcedTaskErrors (black) and StandardTaskErrors (green). Error reduction was slower with NoTaskErrors (red) than with StandardTaskErrors (green), and than with enforced task errors (black). **1b.** Asymptote (mean reach direction in the last 5 adaptation cycles). **1c.** Implicit remapping (mean reach direction in the first no-feedback cycle). Implicit remapping in the NoTaskError group was reduced compared to the StandardTaskError group and the ConstantTaskError group. **1d.** Volitional disengagement of explicit learning, as quantified by the change in reach direction upon notification that the cursor rotation had been removed. **1e.** Group mean+/-SEM for reach directions across all cycles for the follow-up experiment with 30 instead of 60 adaptation cycles. The pattern of slower adaptation with NoTaskErrors in 1a was replicated in this study. **1f.** Less implicit remapping was evident with NoTaskErors than with StandardTaskErrors. **1g.** The NoTaskError group also did not show volitional disengagement of explicit learning. Error bars were standard errors of the mean for cycle by cycle data in **1a** and **1e**. Error bars were 95% confidence intervals for **1b-d** and **1f-h**.

#### Adaptation (cycles 1 to 30)

In the first 30 adaptation cycles, reaches were least adapted with NoTaskErrors (-12.5+/-1.2°), followed by EnforcedTaskErrors (-14.3+/-0.9°), and most adapted with StandardTaskErrors (-24.2+/-0.8°). Cycle x Condition ANOVA showed a significant main effect of Condition, *F*_2,91_ = 36.26, *p* < 0.001, partial η-squared = 0.44, and a Cycle x Condition interaction, *F*_11.7,534.4_ = 3.06, *p* < 0.001, partial η-squared = 0.06. Post-hoc comparisons showed that both the NoTaskError group and the EnforcedTaskError group showed less error compensation compared to StandardTaskErrors (both *p* <.001). Adaptation in the NoTaskError and EnforcedTaskError groups did not differ reliably (*p* = .64).

#### Asymptote (cycles 56 to 60)

To evaluate the completeness of adaptation to the perturbation, we compared asymptotic reach direction, quantified as the mean reach direction in the final 5 cycles (i.e., after more than 400 trials of exposure to the perturbation). The difference between task error conditions remained evident even at asymptote. The EnforcedTaskError and NoTaskError group were not normally distributed due to one outlier in each group (see Fig1b). A Kruskal-Wallis test showed a difference in asymptote between conditions, *chi-square* = 12.7, *p* <.001. Adaptation was least complete with NoTaskErrors (-18.2+/-1.7°), followed by EnforcedTaskErrors (-22.7+/-1.4°) and StandardTaskErrors (-29.7+/-0.6°). StandardTaskErrors resulted in more complete adaptation compared to NoTaskErrors, *U* = 77, *z* = −5.677, *p* < 0.001, *r* = 0.77, as well as compared to EnforcedTaskErrors, *U* = 95, *z* = −5.423, *p* < 0.001, *r* = 0.6. NoTaskErrors resulted in less complete adaptation to the perturbation than EnforcedTaskErrors, *U* = 298, *z* = −2.873, *p* = 0.004, *r* = 0.36.

### Disengagement of explicit learning

Before the first no-feedback cycle, participants were explicitly told that the cursor rotation had been removed: this typically evokes a volitional disengagement of explicit learning, as evident in a change in reach direction between the last adaptation cycle compared to the first no-feedback cycle (Heuer & Hegele, 2008b; a; Hegele & Heuer, 2010). We quantified this volitional disengagement of explicit learning across the different conditions by comparing the last adaptation cycle to the first no-feedback cycle, using a Condition x Cycle (last adaptation cycle, first no-feedback cycle) ANOVA. There was a significant Condition x Cycle interaction, F_2,91_ = 9.69, p < 0.001, partial η-squared = 0.17. Post-hoc paired t-tests comparing the last adaptation cycle to the first no-feedback cycle shows evident disengagement of explicit learning with StandardTaskErrors, *t*_29_= −6.46, p < 0.001, *d* = 1.18, but not with EnforcedTaskErrors, *t*_31_ = −0.78, *p* = 0.44, *d* = 0.13, and not with NoTaskErrors, *t*_31_ = −1.21, *p* = 0.23, *d* = 0.21. The absence of this discrepancy for EnforcedTaskErrors and NoTaskErrors implies that these manipulations of task errors provide a reasonable assay of implicit learning during exposure to the perturbation.

### Removing task error reduced implicit remapping

#### Implicit remapping

Implicit remapping was quantified by the extent to which reach direction remained adapted in the first no-feedback cycle despite knowledge of rotation removal. Similar to previous work (Leow *et al.*, 2017), we only assessed the first no-feedback cycle, as sensorimotor adaptation decays rapidly in the absence of cursor feedback (Kitago *et al.*, 2013).

Figure 1b shows smaller implicit remapping with NoTaskErrors, −18.0+/-1.2°, than StandardTaskErrors, −22.1+/-1.0°, *t*_59_ = 2.63, *p* =.01, cohen’s *d* = 0.67. Remapping was also reduced with NoTaskErrors, −18.0+/-1.2°, compared to ConstantTaskErrors, −21.6+/-0.8, *t*_53.8_ = 2.56, *p* = .013, cohen’s *d* = 0.64. Hence, the absence of task failure in the NoTaskError condition resulted in both less adapted reaches and less post-adaptation implicit remapping than the constant presence of task failure in the EnforcedTaskError condition. Implicit remapping did not differ reliably between StandardTaskErrors and ConstantTaskErrors, *p* >0.5, cohen’s *d* =0.06.

The modest difference (cohen’s *d* =0.67, medium effect size) in implicit remapping between the StandardTaskError and NoTaskError groups might have been due to a ceiling effect resulting from the large number of adaptation cycles. We next assessed data from the follow-up experiment with a different group of naïve participants who encountered 30 instead of 60 adaptation cycles before the no-feedback block, and similarly found less implicit remapping with NoTaskErrors, −15.3+/-0.88°, than StandardTaskErrors, 19.8+/-0.87°, *U* = 54, *z* = −2.789, *p* = 0.005, *r* = −0.70.

## Discussion

Sensorimotor perturbations typically evoke task errors and sensory prediction errors at the same time, making it difficult to disentangle the effects of these distinct error sources on adaptation. In this experiment, we dissociated the effects of task errors from sensory prediction errors during target-reaching by either (1) removing task errors by moving the target to align with the cursor direction mid-movement such that the cursor always hits the target, or (2) enforcing constant task errors by moving the target away from the cursor mid-movement such that the cursor never hits the target, or (3) allowing standard task errors, where the target did not move during the trial. Participants adapted to a 30° rotation of cursor feedback across all conditions. Both removing task errors and enforcing constant task errors reduced the rate and extent of adaptation compared to standard task errors, but this reduction was largest when task errors were removed. After adaptation, we informed participants that the perturbation had been removed: persistently adapted movements despite this knowledge measures indicate an implicit change in the mapping between the motor command and the predicted sensory outcomes of the motor command (i.e., implicit remapping). Removing task errors reduced implicit remapping, whilst constant task errors resulted in similar implicit remapping as standard task errors. This evidence shows that whether or not people attain intrinsic rewards, through hitting/missing targets during goal-directed reaching, makes a crucial contribution to the mapping between motor commands and the predicted sensory outcomes of the motor commands.

### Removing task errors slowed adaptation

We found a dramatic reduction in the rate and extent of adaptation to the perturbation when task errors were removed: during early adaptation, the amount of adaptation was approximately half (-12.5+/-1.2°) of that observed in standard conditions (-24.2+/-0.8°). Previous studies also examined the effects of reducing task errors on adaptation, however, this was accomplished by reducing target precision, for example by having participants aim to an arc-sized target compared to a ray-sized target (Schaefer *et al.*, 2012), or a large target compared to a small target whilst a perturbation was gradually introduced (Reichenthal *et al.*, 2016), or by asking participants to aim where they pleased in the absence of a target (Welch, 1969). Manipulating target errors by altering target precision is known to alter spatial characteristics of movements (Fitts, 1954; Soechting, 1984), possibility due to greater uncertainty about where to aim, and reduced precision of the movement plan (Reichenthal *et al.*, 2016). Nonetheless, despite the differences in methodology, our results are consistent with previous findings: adaptation was slower when target errors were reduced, and reaches were less adapted when the perturbation was removed. Target errors therefore drive the rate and extent of compensation to sensorimotor perturbations.

### Target errors as an intrinsic reward or punishment that contributes to implicit remapping

Participants who never experienced target misses in the NoTaskError condition showed not only reduced extent of adaptation, but also reduced amount of post-adaptation implicit remapping. Hence, removing the negative reward prediction errors associated with target-misses caused by a perturbation reduces implicit remapping to that perturbation. In a similar vein, previous work has demonstrated potent effects of reward on sensorimotor remapping in owls who were either fed dead mice, or hunted live mice during a 10 week exposure to prism glasses. Despite similar feeding durations of 1 hour per day, and similar feeding behaviours (orientation towards the mice, flying, and striking the target), owls who hunted live mice showed a five-fold increase in the shift of auditory space maps in the optic tectum than owls who were fed dead mice (Bergan *et al.*, 2005). We speculate that hunting live mice in the presence of the visual perturbation ensured that the owls had to actively correct for the perturbation in order to attain the food reward. In contrast, the owls who were fed dead mice regardless of their actions were similar to our NoTaskError group, who also showed less remapping than the StandardTaskError group.

Perturbations elicit either an unexpected failure to attain the reward of hitting target (i.e., a negative reward prediction error), and/or an unexpected punishment of missing target (i.e., a positive punishment error). Our results suggest that this reward and/or punishment-based process makes an important contribution to implicit acquisition of new sensorimotor maps in response to sensory prediction errors. This demonstrates an interaction between the system that learns from sensory prediction errors, and the system that learns from reward prediction errors. In contrast, previous theories of adaptation have considered error-based learning and reward-based learning to operate independently from each other, because while sensory prediction errors alters the mapping between motor commands and the predicted sensory consequences of the motor command, reward prediction errors alone do not (Izawa & Shadmehr, 2011; Nikooyan & Ahmed, 2015). This proposal is supported by a large body of work showing that distinct neural systems subserve error-based learning and reward-based learning. Error-based learning is subserved by the cerebellum, as it is required for implicit adaptation of input-output maps between motor commands and sensory outcomes in response to sensory prediction errors (Martin *et al.*, 1996; Werner *et al.*, 2009; Schlerf *et al.*, 2013; Therrien *et al.*, 2015; Butcher *et al.*, 2017). Reward-based learning is known to be subserved by the basal ganglia, which seems likely to be responsible for an action-selection policy that reduces task error (Shadmehr & Krakauer, 2008; Taylor & Ivry, 2014). The finding that the presence or absence of intrinsic rewards associated with hitting or missing targets affects implicit adaptation to sensory prediction errors suggests an alternative hypothesis: that error-based learning subserved by the cerebellum is also sensitive to reward/punishment signals from the basal ganglia. This seems plausible given the presence of disynaptic projections between the cerebellum and the basal ganglia (Bostan, Dum et al. 2010, Bostan and Strick 2010), which might allow reward signals processed by the basal ganglia to modulate implicit adaptation driven by sensory prediction errors in the cerebellum. New evidence also shows that the cerebellum is sensitive to rewards (Herzfeld *et al.*, 2015). For example, cerebellar granule cells encode reward expectation, as their activity peaks in the pre-reward period (Wagner *et al.*, 2017). The post-synaptic targets of cerebellar granule cells are Purkinje cells, which are known to play a crucial role in sensorimotor adaptation (Herzfeld *et al.*, 2015).

Neuropsychological evidence in humans supports the hypothesis that implicit acquisition of a new sensorimotor map is sensitive to reward. For example, when the neurotransmitter required for processing reward, dopamine, is deficient in Parkinson’s disease, post-adaptation aftereffects are also reduced (Stern *et al.*, 1988; Contreras-Vidal & Buch, 2003; Fernandez-Ruiz *et al.*, 2003; Gutierrez-Garralda *et al.*, 2013; Roemmich *et al.*, 2014). Withdrawing dopamine medication in Parkinson’s disease patients further reduces the size of the aftereffect (Roemmich *et al.*, 2014), demonstrating a role for dopamine reward signals in implicit remapping. On the other hand, if reward prediction errors typically modulate processing of sensory prediction errors by the cerebellum, then impaired cerebellar function might not only impair the capacity for implicit adaptation to sensory prediction errors, but might also impair the capacity to respond appropriately to reward prediction errors. This is reflected in the apparent deficit cerebellar degeneration patients in independently developing a strategy and re-aiming in response to reward prediction errors (Therrien *et al.*, 2015; Butcher *et al.*, 2017), despite intact ability to implement a strategy provided by the experimenter (Taylor *et al.*, 2010).

### Enforced task errors slowed error compensation

We enforced task errors by moving the target away from the cursor randomly clockwise or counterclockwise by 20° to 30°, such that participants could never hit the target regardless of how they moved. This manipulation is similar to previous work which clamps cursor feedback to a constant offset away from the target, regardless of how participants moved (Kim *et al.*, 2017a; Morehead *et al.*, 2017), although those studies actually intended to remove task error by instructing participants to ignore the clamped cursor feedback. If participants succeed at obeying instructions to ignore the feedback, they technically do not commit any task errors (Welch, 1969). We suggest however that clamped cursor feedback actually enforces constant target errors, because participants observe their cursor constantly failing to hit the target, regardless of where they reach. These studies showed slower adaptation with clamped cursor feedback than with standard cursor feedback, but implicit remapping did not differ with standard or clamped cursor feedback (Kim *et al.*, 2017a; Morehead *et al.*, 2017). Similarly, adaptation was slower with our constant task error group than our standard task error group, and implicit remapping did not differ between standard or constant task error conditions.

A strength of the current work is that we compared how removing task errors and enforcing task errors affected adaptation, unlike previous work which examined these in isolation. Removing task errors removes the motivation for strategy use, whereas enforcing constant task errors deters strategy use by ensuring that all strategies are futile. Both methods appeared successful in suppressing strategy use: participants in both conditions showed no change in behaviour before and after explicit knowledge of perturbation removal. Despite this, implicit remapping was smaller with NoTaskErrors than with EnforcedTaskErrors: strategy use thus might not underlie this difference in implicit remapping. Furthermore, implicit remapping resulting from EnforcedTaskErrors did not differ reliably from StandardTaskErrors. We speculate that even when task errors cannot provide a strategy for task success, task errors remain an important component to the process of implicit remapping.

### Intrinsic versus extrinsic rewards and punishments

We operationalized target hits as intrinsically rewarding and target misses to be intrinsically punishing, without providing any additional extrinsic rewards or punishments such as monetary gains or monetary losses. We do not know if and how extrinsic rewards or punishments might interact with intrinsic rewards or punishments during learning. The majority of previous work examining the role of reward on motor learning manipulated extrinsic rewards and punishments without manipulating intrinsic rewards and punishments (e.g., Wachter *et al.*, 2009; Abe *et al.*, 2011; Galea *et al.*, 2015; Nikooyan & Ahmed, 2015; Gajda *et al.*, 2016; Steel *et al.*, 2016; Song & Smiley-Oyen, 2017). One possibility is that extrinsic rewards and punishments affect learning via additive or subtractive effects on intrinsic reward processes. This is because extrinsic rewards and punishments only affected learning when they were meted out in conjunction with task errors: providing rewards or punishments that were not contingent upon task errors did not alter learning (Galea *et al.*, 2015; Nikooyan & Ahmed, 2015). Alternatively, extrinsic and intrinsic rewards might exert independent effects on learning. These possibilities await future study.

### Summary

In summary, we showed that the intrinsic reward of hitting or missing targets during target-reaching in sensorimotor adaptation affects implicit remapping resulting from exposure to sensory prediction errors. Hence, even though reward prediction errors alone are not sufficient to result in sensory remapping (Izawa & Shadmehr, 2011), reward prediction errors appear to contribute to changes in sensory remapping, possibly by increasing sensitivity to sensory prediction errors.

## References

Abe, M., Schambra, H., Wassermann, E.M., Luckenbaugh, D., Schweighofer, N. & Cohen, L.G. (2011) Reward improves long-term retention of a motor memory through induction of offline memory gains. Current Biology, 21, 557–562.

Bergan, J.F., Ro, P., Ro, D. & Knudsen, E.I. (2005) Hunting increases adaptive auditory map plasticity in adult barn owls. J Neurosci, 25, 9816–9820.

Butcher, P.A., Ivry, R., Kuo, S.-H., Rydz, D., Krakauer, J.W. & Taylor, J.A. (2017) The Cerebellum Does More Than Sensory-Prediction-Error-Based Learning In Sensorimotor Adaptation Tasks. bioRxiv, 139337.

Cashaback, J.G., McGregor, H.R., Mohatarem, A. & Gribble, P.L. (2017) Dissociating error-based and reinforcement-based loss functions during sensorimotor learning. PLoS computational biology, 13, e1005623.

Contreras-Vidal, J.L. & Buch, E.R. (2003) Effects of Parkinson’s disease on visuomotor adaptation. Experimental Brain Research, 150, 25–32.

Fernandez-Ruiz, J., Diaz, R., Hall-Haro, C., Vergara, P., Mischner, J., Nunez, L., Drucker-Colin, R., Ochoa, A. & Alonso, M.E. (2003) Normal prism adaptation but reduced after-effect in basal ganglia disorders using a throwing task. European Journal of Neuroscience, 18, 689–694.

Fitts, P.M. (1954) The information capacity of the human motor system in controlling the amplitude of movement. Journal of Experimental Psychology, 47, 381–391.

Fritz, C.O., Morris, P.E. & Richler, J.J. (2012) Effect size estimates: current use, calculations, and interpretation. Journal of experimental psychology: General, 141, 2.

Gajda, K., Sülzenbrück, S. & Heuer, H. (2016) Financial incentives enhance adaptation to a sensorimotor transformation. Experimental Brain Research, 234, 2859–2868.

Galea, J.M., Mallia, E., Rothwell, J. & Diedrichsen, J. (2015) The dissociable effects of punishment and reward on motor learning. Nat Neurosci, 18, 597–602.

Ghilardi, M.F., Gordon, J. & Ghez, C. (1995) Learning a visuomotor transformation in a local area of work space produces directional biases in other areas. Journal of Neurophysiology, 73, 2535–2539.

Gutierrez-Garralda, J.M., Moreno-Briseño, P., Boll, M.C., Morgado-Valle, C., Campos-Romo, A., Diaz, R. & Fernandez-Ruiz, J. (2013) The effect of Parkinson’s disease and Huntington’s disease on human visuomotor learning. European Journal of Neuroscience, 38, 2933–2940.

Hegele, M. & Heuer, H. (2010) Implicit and explicit components of dual adaptation to visuomotor rotations. Consciousness and Cognition, 19, 906–917.

Herzfeld, D.J., Kojima, Y., Soetedjo, R. & Shadmehr, R. (2015) Encoding of action by the Purkinje cells of the cerebellum. Nature, 526, 439–442.

Heuer, H. & Hegele, M. (2008a) Adaptation to visuomotor rotations in younger and older adults. Psychol Aging, 23, 190–202.

Heuer, H. & Hegele, M. (2008b) Constraints on visuo-motor adaptation depend on the type of visual feedback during practice. Experimental Brain Research, 185, 101–110.

Howard, I.S., Ingram, J.N. & Wolpert, D.M. (2009) A modular planar robotic manipulandum with end-point torque control. Journal of Neuroscience Methods, 181, 199–211.

Izawa, J. & Shadmehr, R. (2011) Learning from sensory and reward prediction errors during motor adaptation. PLoS Comput Biol, 7, e1002012.

Kim, H., Morehead, J.R., Parvin, D., Moazzezi, R. & Ivry, R. (2017a) Invariant errors reveal limitations in motor correction rather than constraints on error sensitivity. bioRxiv, 189597.

Kim, H.E., Parvin, D.E., Hernandez, M.A. & Ivry, R.B. (2017b) Implicit rewards modulate sensorimotor adaptation Advances in Motor Learning and Motor Control (MLMC), San Diego.

Kitago, T., Ryan, S.L., Mazzoni, P., Krakauer, J.W. & Haith, A.M. (2013) Unlearning versus savings in visuomotor adaptation: Comparing effects of washout, passage of time and removal of errors on motor memory. Frontiers in Human Neuroscience.

Kojima, Y. & Soetedjo, R. (2017) Selective reward affects the rate of saccade adaptation. Neuroscience, 355, 113–125.

Leow, L.A., Gunn, R., Marinovic, W. & Carroll, T.J. (2017) Estimating the implicit component of visuomotor rotation learning by constraining movement preparation time. J Neurophysiol, jn 00834 02016.

Madelain, L., Paeye, C. & Wallman, J. (2011) Modification of saccadic gain by reinforcement. Journal of neurophysiology, 106, 219–232.

Martin, T.A., Keating, J.G., Goodkin, H.P., Bastian, A.J. & Thach, W.T. (1996) Throwing while looking through prisms I. Focal olivocerebellar lesions impair adaptation. Brain, 119, 1183–1198.

Mazzoni, P. & Krakauer, J.W. (2006) An implicit plan overrides an explicit strategy during visuomotor adaptation. Journal of Neuroscience, 26, 3642–3645.

McDougle, S.D., Ivry, R.B. & Taylor, J.A. (2016) Taking aim at the cognitive side of learning in sensorimotor adaptation tasks. Trends in cognitive sciences, 20, 535–544.

Morehead, J.R. & Ivry, R. (2015) Intrinsic biases systematically affect visuomotor adaptation experiments. Neural Control of Movement, Charleston.

Morehead, J.R., Taylor, J.A., Parvin, D.E. & Ivry, R.B. (2017) Characteristics of Implicit Sensorimotor Adaptation Revealed by Task-irrelevant Clamped Feedback. Journal of Cognitive Neuroscience.

Nikooyan, A.A. & Ahmed, A.A. (2015) Reward feedback accelerates motor learning. J Neurophysiol, 113, 633–646.

Palidis, D.J., Cashaback, J. & Gribble, P. (2018) Distinct Neural Signatures of Reward and Sensory Prediction Error in Motor Learning. bioRxiv, 262576.

Quattrocchi, G., Greenwood, R., Rothwell, J.C., Galea, J.M. & Bestmann, S. (2017) Reward and punishment enhance motor adaptation in stroke. Journal of Neurology, Neurosurgery and Psychiatry, 88, 730–736.

Reichenthal, M., Avraham, G., Karniel, A. & Shmuelof, L. (2016) Target size matters: Target errors contribute to the generalization of implicit visuomotor learning. Journal of neurophysiology, jn. 00830.02015.

Roemmich, R.T., Hack, N., Akbar, U. & Hass, C.J. (2014) Effects of dopaminergic therapy on locomotor adaptation and adaptive learning in persons with Parkinson’s disease. Behavioural Brain Research, 268, 31–39.

Schaefer, S.Y., Shelly, I.L. & Thoroughman, K.A. (2012) Beside the point: Motor adaptation without feedback-based error correction in task-irrelevant conditions. Journal of Neurophysiology, 107, 1247–1256.

Schlerf, J.E., Xu, J., Klemfuss, N.M., Griffiths, T.L. & Ivry, R.B. (2013) Individuals with cerebellar degeneration show similar adaptation deficits with large and small visuomotor errors. J Neurophysiol, 109, 1164–1173.

Schouten, J.F. & Bekker, J.A.M. (1967) Reaction time and accuracy. Acta Psychologica, 27, 143–153.

Shadmehr, R. & Krakauer, J.W. (2008) A computational neuroanatomy for motor control. Experimental Brain Research, 185, 359–381.

Soechting, J. (1984) Effect of target size on spatial and temporal characteristics of a pointing movement in man. Experimental Brain Research, 54, 121–132.

Song, Y. & Smiley-Oyen, A.L. (2017) Probability differently modulating the effects of reward and punishment on visuomotor adaptation. Experimental Brain Research, 1–14.

Steel, A., Silson, E.H., Stagg, C.J. & Baker, C.I. (2016) The impact of reward and punishment on skill learning depends on task demands. Scientific reports, 6.

Stern, Y., Mayeux, R., Hermann, A. & Rosen, J. (1988) Prism adaptation in Parkinson’s disease. J Neurol Neurosurg Psychiatry, 51, 1584–1587.

Taylor, J.A. & Ivry, R.B. (2011) Flexible cognitive strategies during motor learning. Plos Computational Biology, 7.

Taylor, J.A. & Ivry, R.B. (2014) Cerebellar and Prefrontal Cortex Contributions to Adaptation, Strategies, and Reinforcement Learning Progress in Brain Research, pp. 217–253.

Taylor, J.A., Klemfuss, N.M. & Ivry, R.B. (2010) An Explicit Strategy Prevails When the Cerebellum Fails to Compute Movement Errors. Cerebellum, 9, 580–586.

Therrien, A.S., Wolpert, D.M. & Bastian, A.J. (2015) Effective reinforcement learning following cerebellar damage requires a balance between exploration and motor noise. Brain, awv329.

van der Kooij, K. & Overvliet, K.E. (2016) Rewarding imperfect motor performance reduces adaptive changes. Experimental Brain Research, 234, 1441–1450.

Vindras, P. & Viviani, P. (1998) Frames of Reference and Control Parameters in Visuomanual Pointing. Journal of Experimental Psychology: Human Perception and Performance, 24, 569–591.

Wachter, T., Lungu, O.V., Liu, T., Willingham, D.T. & Ashe, J. (2009) Differential Effect of Reward and Punishment on Procedural Learning. Journal of Neuroscience, 29, 436–443.

Wagner, M.J., Kim, T.H., Savall, J., Schnitzer, M.J. & Luo, L. (2017) Cerebellar granule cells encode the expectation of reward. Nature, 544, 96–100.

Welch, R.B. (1969) Adaptation to prism-displaced vision: The importance of target-pointing. Perception & Psychophysics, 5, 305–309.

Werner, S., Bock, O. & Timmann, D. (2009) The effect of cerebellar cortical degeneration on adaptive plasticity and movement control. Exp Brain Res, 193, 189–196.

Widmer, M., Ziegler, N., Held, J., Luft, A. & Lutz, K. (2016) Rewarding feedback promotes motor skill consolidation via striatal activity. Elsevier B.V.

